# Predicting cognitive and mental health traits and their polygenic architecture using large-scale brain connectomics

**DOI:** 10.1101/609586

**Authors:** Luigi A. Maglanoc, Tobias Kaufmann, Dennis van der Meer, Andre F. Marquand, Thomas Wolfers, Rune Jonassen, Eva Hilland, Ole A. Andreassen, Nils Inge Landrø, Lars T. Westlye

**Affiliations:** Clinical Neuroscience Research Group, Department of Psychology, University of Oslo, Oslo, Norway; NORMENT, Division of Mental Health and Addiction, Oslo University Hospital & Institute of Clinical Medicine, University of Oslo, Norway; School of Mental Health and Neuroscience, Faculty of Health, Medicine and Life Sciences, Maastricht University, The Netherlands; Donders Centre for Cognitive Neuroimaging, Donders Institute for Brain, Cognition and Behaviour, Radboud University, Nijmegen, The Netherlands; Department of Neuroimaging, Institute of Psychiatry, King’s College London, London, UK; Department of Health Sciences, Oslo Metropolitan University, Oslo, Norway; Division of Psychiatry, Diakonhjemmet Hospital, Oslo, Norway; Department of Psychology, University of Oslo, Oslo, Norway

**Keywords:** fMRI, functional connectivity, human traits, polygenic scores, machine learning

## Abstract

Cognitive abilities and mental disorders are complex traits sharing a largely unknown neuronal basis and aetiology. Their genetic architectures are highly polygenic and overlapping, which is supported by heterogeneous phenotypic expression and substantial clinical overlap. Brain network analysis provides a non-invasive means of dissecting biological heterogeneity yet its sensitivity, specificity and validity in clinical applications remains a major challenge. We used machine learning on static and dynamic temporal synchronization between all brain network nodes in 10,343 healthy individuals from the UK Biobank to predict (i) cognitive and mental health traits and (ii) their genetic underpinnings. We predicted age and sex to serve as our reference point. The traits of interest included individual level educational attainment and fluid intelligence (cognitive) and dimensional measures of depression, anxiety, and neuroticism (mental health). We predicted polygenic scores for educational attainment, fluid intelligence, depression, anxiety, and different neuroticism traits, in addition to schizophrenia. Beyond high accuracy for age and sex, permutation tests revealed above chance-level prediction accuracy for educational attainment and fluid intelligence. Educational attainment and fluid intelligence were mainly negatively associated with static brain connectivity in frontal and default mode networks, whereas age showed positive correlations with a more widespread pattern. In comparison, prediction accuracy for polygenic scores was at chance level across traits, which may serve as a benchmark for future studies aiming to link genetic factors and fMRI-based brain connectomics.

**Significance:** Although cognitive abilities and susceptibility to mental disorders reflect individual differences in brain function, neuroimaging is yet to provide a coherent account of the neuronal underpinnings. Here, we aimed to map the brain functional connectome of (i) cognitive and mental health traits and (ii) their polygenic architecture in a large population-based sample. We discovered high prediction accuracy for age and sex, and above-chance accuracy for educational attainment and intelligence (cognitive). In contrast, accuracies for dimensional measures of depression, anxiety and neuroticism (mental health), and polygenic scores across traits, were at chance level. These findings support the link between cognitive abilities and brain connectomics and provide a reference for studies mapping the brain connectomics of mental disorders and their genetic architectures.

## Introduction

Mental health and cognitive abilities are both linked to brain function. Mental disorders such as depression and schizophrenia have joint high prevalence, early onset, and often a persistent nature (1). As such, identifying neuroimaging-based biomarkers which may facilitate early risk detection and interventions is a global aim potentially implicating millions worldwide (2–4). Twin and family studies have documented a strong genetic contribution across a range of human traits and mental disorders (5), and high polygenicity based on genome-wide associations studies (GWAS)(6–10). Educational attainment and fluid intelligence are two related and heritable phenotypes associated with a range of behaviors and outcomes (11) such as occupational attainment and social mobility (12), brain measures (13–15), and health (16, 17). Further, they show strong genetic correlations across cognitive domains and levels of ability (18).

Both the structural and functional architecture of the brain are heritable traits (19), and previous studies reporting aberrations in fMRI measures of brain connectivity in mental disorders are abundant (20–22). However, the generalizability, robustness and clinical utility of the findings have been questioned (23–25). This is likely due to differences in power (26), sample characteristics and analytical methods (27), but also the vast clinical heterogeneity of mental disorders (28–30). Further, although a substantial heritability of brain measures has been demonstrated (19, 31, 32), the sensitivity of fMRI features to the polygenic risk for mental disorders and associated personality traits is largely unknown.

Studies assessing the heritability and clinical associations with fMRI resting-state brain functional connectivity (FC) have typically targeted estimates of static FC (sFC), defined as the average temporal correlation between two brain regions. Less is known about the associations with dynamic properties of FC (dFC), conceptualized as the fluctuation in temporal correlations between two brain regions. Further, fMRI-based FC reflects the joint contribution of partly independent sources oscillating at specific frequency bands (33), which are tied to a range of neural processes (34) and cognitive functions (35). Establishing the sensitivity and differential associations of imaging-based indices of brain function and connectivity to the genetic architecture of cognitive abilities and mental disorders may inform nosological and mechanistic studies and improve diagnostics, prevention and treatment.

Here, we tested the ability to detect mappings between static and dynamic measures of brain connectivity with (i) cognitive and mental health traits and (ii) genetic architecture of related traits based on polygenic score. These were compared with mappings of age and sex to help establish the results as a benchmark for clinical functional imaging We used resting-state fMRI data from 10,343 healthy individuals from the UK Biobank (UKB) in a multivariate machine learning approach. The cognitive traits were years of educational attainment and fluid intelligence, and the mental health traits were dimensional measures of depression, anxiety and neuroticism. The polygenic scores in the same individuals included educational attainment, fluid intelligence, depression, anxiety, and 13 neuroticism traits. In addition, we included polygenic scores for schizophrenia, which is a disorder with high relevance for brain function, but currently not possible to study directly in the UKB due to the low number of cases with available MRI data. We employed cross-validation and evaluation of model performance to reduce bias and overfitting, in addition to permutation testing for statistical inference.

## Results

### Multivariate prediction of phenotypes

Fig. 1 shows the distribution of the phenotypes and cross-validated results of the overall best feature set (sFC) using 10-fold internal cross-validation with 100 repetitions (on 80% of the total sample). Permutation testing revealed above chance-level prediction accuracy for phenotypic level educational attainment, fluid intelligence, age, (see Table 1) and sex (accuracy = 77.3%, corrected *p* < 0.0027, sensitivity = 87.7%, specificity = 32.4%). In comparison, predictions of dimensional measures of depression, anxiety and neuroticism anxiety performed at chance level (see Table 1). Fig. S1 and S2 provides the cross-validated results of the other FC feature sets. The feature set with all three FC-types (sFC, bandpass filtered sFC, and dFC) resulted in marginally higher prediction accuracy than the sFC feature set only for age (*t* = 85.613, df = 170.83, *p* < 0.001) and sex (*t* = 75.785, df = 183.04, *p* < 0.001). The model validation results (using 80% of the total sample to predict the leftover 20%) are very similar to the cross-validated results (see Fig. S3). Sensitivity analyses predicting age using the sFC feature set showed that there was very little difference when regressing scanner site out of the edges or excluding participants scanned in Newcastle (See methods and Fig. S4A). Further sensitivity analyses predicting fluid intelligence using sFC showed similar results when regressing out age (linear and quadratic), sex and head motion from the edges in the sFC feature set (see Fig. S4B).

**Fig. 1.**
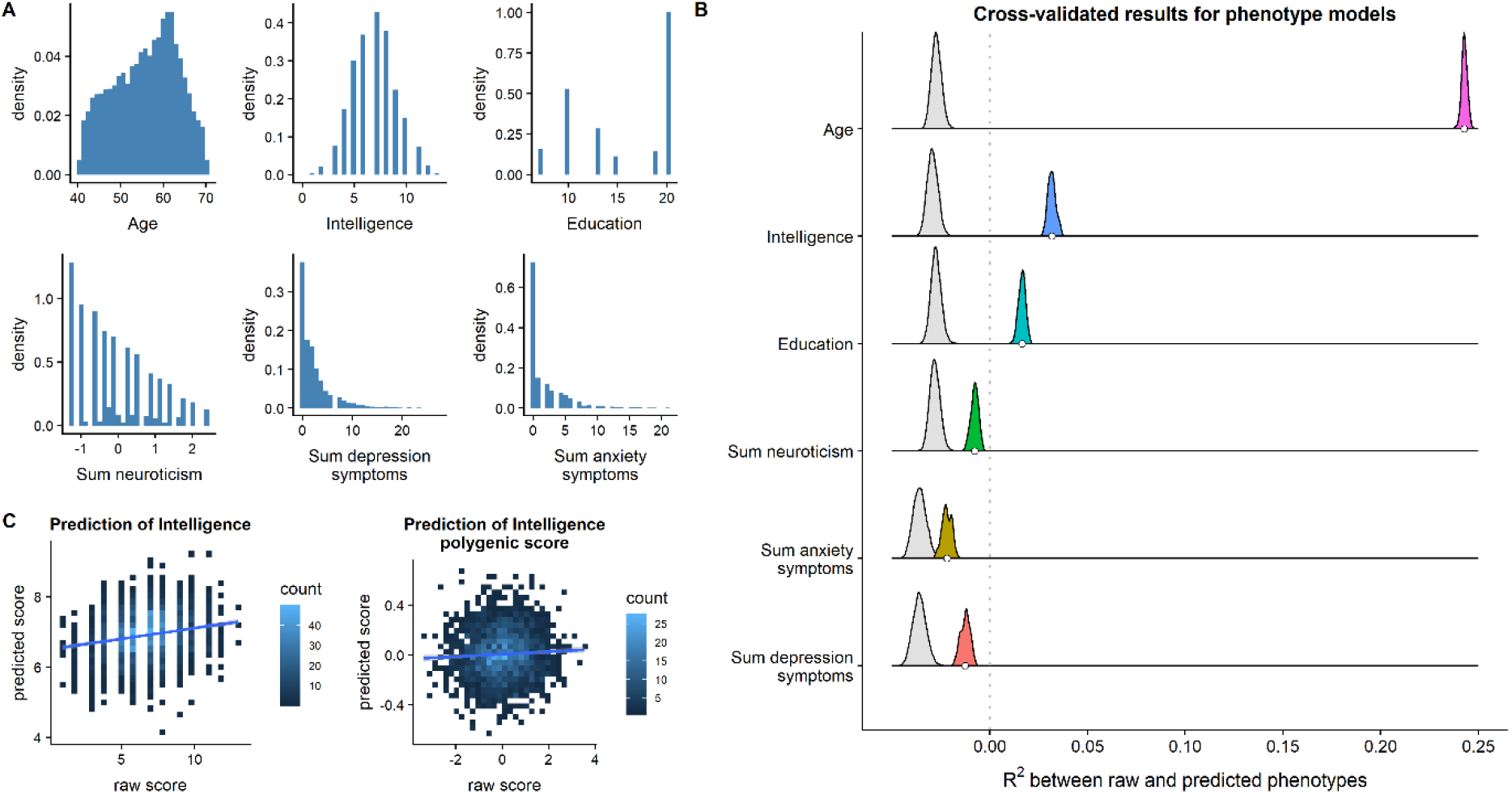
(A) The distribution of all the phenotypes we used in the multivariate analyses. (B) The coloured density distributions represent the cross-validated results (across 100 repetitions) of the best FC feature set (sFC: the mean is denoted by the white circles) for a given phenotype, using R^2^ as our metric of model performance. The grey distributions represent the mean model performance for each of the 10,000 permuted datasets. Although RMSE is our main metric of model performance, R^2^ allows for comparison across phenotypes and polygenic risk scores. R^2^ is negative for some of the models because we are testing it out-of-sample, meaning that the model can be arbitrarily bad. (C) Associations between raw and predicted scores for phenotypic level intelligence (left) and polygenic score (right). Education pertains to the years of educational attainment

**Table 1:** Model performance measures for all the phenotypes and polygenic scores with high correlation-based model performance using the sFC feature set. Worry is one of the 13 neuroticism traits. Correlation was based on spearman’s rho for the phenotypes and Pearson’s r for the polygenic scores. The *p*-value is calculated using permutation testing (based on RMSE), with an additional Bonferroni correction for 7 phenotypes and 20 polygenic score models.

| | RMSE | *p*-value | Correlation | MAE | $R^2$ |
| --- | --- | --- | --- | --- | --- |
| Cognitive and mental health trait |  |  |  |  |  |
| Educational attainment | 4.739 | < 0.0027 | 0.1634 | 4.348 | 0.0164 |
| Fluid intelligence | 2.043 | < 0.0027 | 0.2014 | 1.638 | 0.0317 |
| Age | 6.482 | < 0.0027 | 0.4947 | 5.317 | 0.2428 |
| Neuroticism | 0.984 | $\geq 1$ | 0.0924 | 0.8201 | -0.0078 |
| Depression | 3.383 | $\geq 1$ | 0.0947 | 2.353 | -0.0126 |
| Anxiety | 3.222 | $\geq 1$ | 0.0616 | 2.270 | -0.0220 |
| Polygenic scores |  |  |  |  |  |
| Educational attainment | 0.997 | $\geq 1$ | 0.0941 | 0.7926 | -0.0061 |
| Fluid intelligence | 1.002 | $\geq 1$ | 0.0553 | 0.8009 | -0.0163 |
| Worry | 1.007 | $\geq 1$ | 0.0670 | 0.8170 | -0.0138 |

Fig. 2 shows the grouping of brain networks based on hierarchical clustering and the top 20 edges in the sFC feature set for trait-level educational attainment and fluid intelligence based on feature importance, using correlation-adjusted marginal correlation (CAR) scores. Briefly, we observed mainly negative association with sFC for educational attainment, particularly within the cluster of default mode network (DMN) and frontal network nodes, and a similar brain functional pattern for fluid intelligence. In contrast, the CAR scores revealed largely positive associations between sFC and age, mainly within the cluster of motor/somatosensory and attentional networks, and within the cluster of default mode and frontal networks. Fig. S5 shows the top 40 edges that were positively associated with each phenotype, based on the feature set using all three FC-types, suggesting that these are largely non-overlapping. In short, based on feature importance, we observed roughly equal amounts of edges across all three FC-types for educational attainment. We observed only two edges based on bandpass filtered sFC between visual networks and the cluster of default mode and frontal networks for fluid intelligence and only three edges based on dFC for age. Table S1 shows the correlation between the FC-types for age, fluid intelligence and educational attainment.

**Fig. 2.**
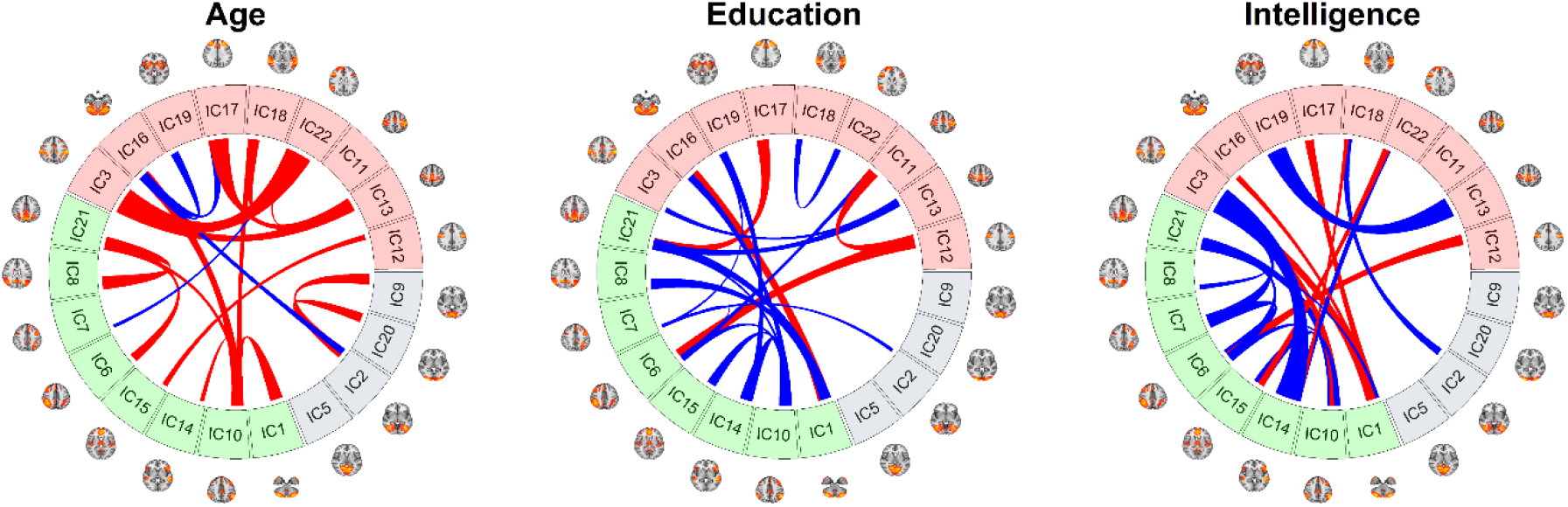
The top 20 edges based on CAR-scores for the best FC feature set (sFC). Cross-fold validation for age, phenotypic years of educational attainment and fluid intelligence. The thickness of the lines represents the relative importance of each edge for that specific feature set. Red lines indicate that the feature is positively associated with the phenotype when considering all the features. Blue lines indicate that the feature is negatively associated with the phenotype when considering all the features. The brain networks are grouped into 3 clusters which roughly represent the motor/somatosensory and attentional networks (red), default mode and frontal networks (green), visual components (blue).

### Multivariate prediction of polygenic scores

Fig. S6 shows the distribution of each polygenic score, and Fig. S7 shows a correlation plot with a dendrogram based on hierarchical clustering across all polygenic scores. Model performance for all polygenic scores (depression, anxiety, schizophrenia, neuroticism traits, educational attainment and intelligence) had negative explained variance based on the coefficient of determination (R^2^) across feature sets (Fig. S8). As such, we only performed permutations testing for the polygenic scores that showed higher correlation-based model performance, including educational attainment, fluid intelligence and worry (Table 1). Fig. 3 provides an illustrative example of relative high model performance (phenotypic level educational attainment) and low model performance (educational attainment polygenic scores) showing both cross-validated results on empirical and permuted data. The model validation results are very similar to the cross-validated results (Fig. S9). There was very little difference when controlling for population stratification by regressing out the first ten genetic principal components for the sFC feature set predicting educational attainment polygenic risk scores (Fig. S4C). Cross-validated results were very similar using a lower polygenic score threshold (p ≤ 0.05: Fig. S10), and after performing principal component analysis (PCA) across p-value thresholds on each of the above individual polygenic scores (36) (Fig. S11). See Supplemental Results for the correlation between phenotypes for specific feature sets, and phenotypes with their corresponding polygenic scores.

**Fig. 3.**
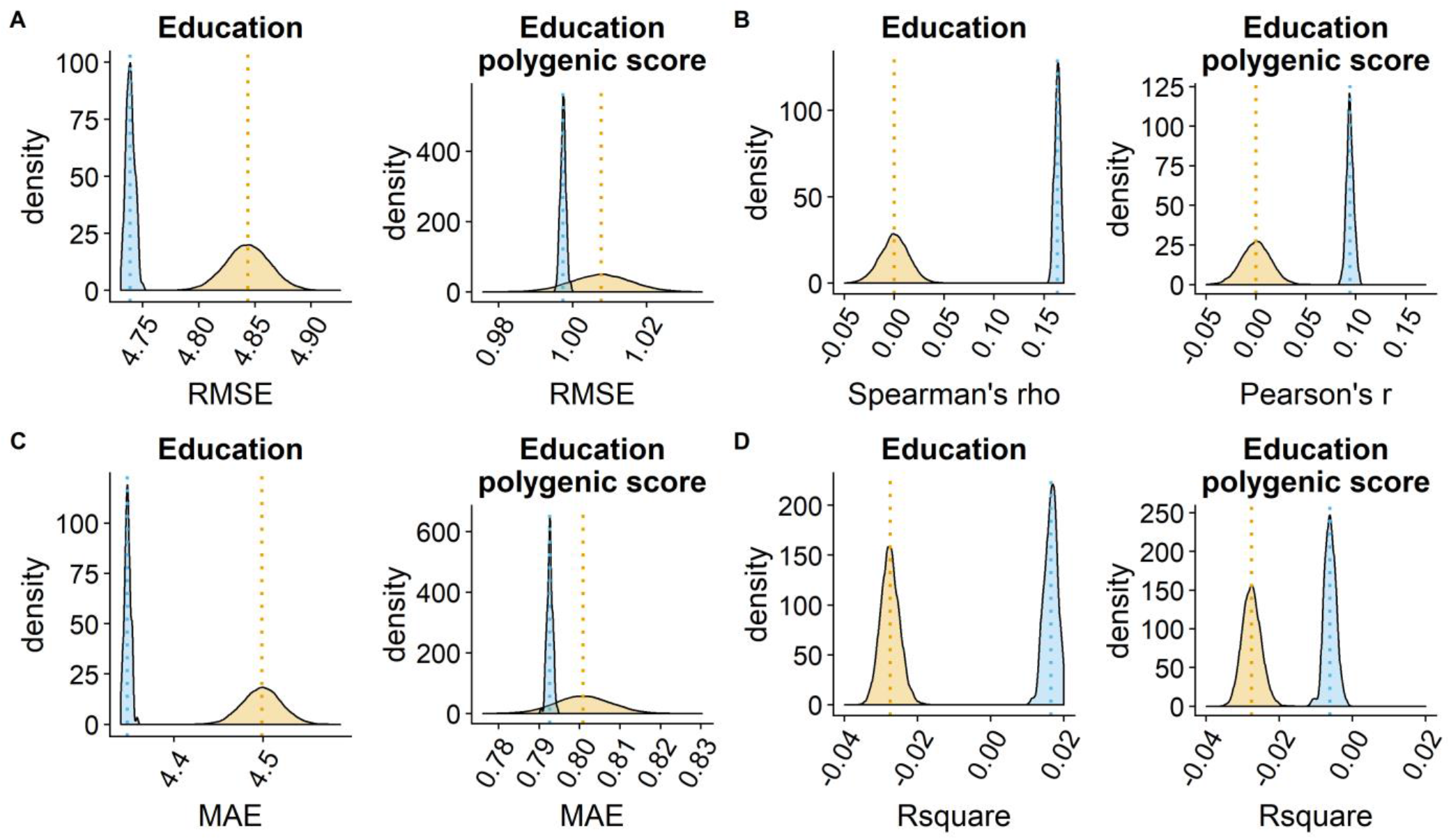
Cross-validated results comparing phenotypic level years of educational attainment and polygenic scores. For each plot, the blue represents model performance for each repetition (100) of the empirical data, and orange represents the mean model performance for each of the 10,000 permuted datasets (orange) based on (A) RMSE, (B) Correlation, (C) mean absolute error (MAE), and (D) R^2^. The lines denote the mean of the respective distributions.

## Discussion

Recent advances have led to an increasing interest in mapping the brain connectomic landscape underlying (i) cognitive and mental health traits and (ii) their genetic underpinnings. To this end, we conducted as large-scale integration of advanced measures of fMRI-based brain functional connectivity in a population cohort of 10,343 healthy individuals. First, we show that individual differences in educational attainment and fluid intelligence (cognitive), which have high relevance for life satisfaction and real-world outcomes, are partly reflected in the temporal organization of the brain. Furthermore, we demonstrate high prediction accuracy for age and sex, which provides a reference point for brain connectomic mapping. In contrast, we found chance-level performance for individual level prediction of dimensional measures of depression, anxiety and neuroticism (mental health). Second, we found chance-level performance for polygenic scores of depression, anxiety, schizophrenia, neuroticism traits, educational attainment and intelligence based on large-scale GWAS.

We found significant prediction accuracy for educational attainment and the corresponding high feature importance for the DMN and frontal networks. We also obtained significant prediction accuracy of brain FC patterns for fluid intelligence, largely driven by negative correlations between intelligence and sFC amongst frontal and DMN networks. This indicates relative dedifferentiation between the DMN and frontal regions in individuals with higher intelligence. This pattern was largely overlapping with the results for educational attainment, consistent with a study using a subset of the current sample (14), and also reflected in a strong positive correlations between the estimated importance of the feature sets from the two prediction tasks.

Highest model performance across all traits was observed for age, in line with several studies showing high age-sensitivity of functional imaging features (37–40). In general, compared to educational attainment and intelligence, we observed a more distributed pattern of the edge-wise feature importance scores for the age prediction. This suggests that individual differences in cognitive abilities may be more confined to specific parts of the brain connectome than the effects of aging, which are more pervasive.

To date, there is a lack of functional imaging studies comparing the sensitivity and predictive accuracy of dynamic and static measures of brain connectivity. Our comparisons revealed similar relative trait ranking in prediction accuracies for the different edge conceptualizations. However, corresponding CAR scores revealed differential feature importance rankings, suggesting that the different edge definitions may capture partly nonoverlapping variance. For age, whereas previous studies have shown that dFC decreases with aging (41, 42), we obtained noticeably poorer prediction accuracy for dFC than the other feature sets, implying that age is better characterized by average FC rather than dynamic FC.

Overall, our results suggest higher dFC within a frontal-DMN cluster in individuals with higher intelligence and educational attainment. However, we must be careful in over-interpreting the dFC and bandpass filtered sFC results, as they performed worse than the feature set with only sFC edges. One exception was that the feature set with all FC-types performed marginally better when predicting age and classifying sex compared to sFC alone, which may suggest that there is added predictive ability when the signal is strong. A recent study found that the discriminative properties of FC vary across parcellation schemes and frequencies, with ICA-based parcellations revealing greater discriminability at high frequencies compared to other parcellations (43).

We were not able to predict dimensional measures of depression or anxiety, consistent with a smaller independent study (44) and a recent large-scale multi-site study (45), or trait neuroticism. One explanation for the lack of brain FC associations, particularly with regards to depression is its diversity in symptom profiles (28), and phenomenological overlap with other clinical domains (46). Some studies have attempted to identify putative subgroups of patients by using data-driven clustering based on symptoms (44, 47) or brain functional connectivity patterns (48, 49). In line with recent work questioning the robustness and generalizability of such clustering approaches (23), our current results do not support a simple link between fMRI-based brain connectivity and sub-clinical manifestations of mental disorders.

To the best of our knowledge, this is the first study that uses a machine learning approach to predict polygenic scores based on resting-state brain FC measures. A recent study demonstrated significant prediction accuracy for polygenic scores for autism using grey matter volumes, but not for attention deficit-hyperactivity disorder, bipolar disorder or schizophrenia (50). However, the sample size was small compared to the current study. Large sample sizes are needed to establish reliable estimates for the neuronal correlates of measures with weak associations (51). Relatedly, the current findings illustrate that the correlation between raw and predicted scores may overestimate model performance, at least when model performance is relatively low. Unlike R^2^, MAE and RMSE, correlation coefficients are affected by scaling and location of the predicted scores relative to the raw scores, and, in contrast to MAE and RMSE, assumes linearity and homoscedasticity. As a result, future studies using multivariate analyses to predict continuous measures should carefully consider the choice of metric used for evaluating model performance, and reporting converging evidence of model fit and robustness may increase the reliability of the findings.

Despite the overall low predictive accuracy of the polygenic scores, we observed a relatively high positive correlation between the edge-wise feature importance of trait level educational attainment and intelligence with their respective polygenic scores, indicating some shared signal. It is conceivable that any genetic pleiotropy between complex human traits and the brain is stronger for other imaging modalities. Indeed, some studies have demonstrated an association between cortical structure and polygenic scores for schizophrenia (52, 53), which are supported by a recent large-scale UKB study reporting thinner fronto-temporal cortex with higher polygenic risk (36), suggesting shared mechanisms. However, the variance explained by polygenic scores, such as those for schizophrenia, in brain measures like gray matter volume, tend to be low, around a few percent (54).

The low predictive accuracy for polygenic scores was observed both when applying a liberal and a more conservative initial p-value threshold when computing the polygenic score, and when employing a PCA-based approach calculated across a wide range of thresholds. Hence, the low predictive accuracy cannot simply be explained by the number of independent single nucleotide polymorphisms (SNP) feeding into the cumulative score. Relatedly, the poor predictive accuracy may also be partly explained by the low amount of explained trait variance accounted for by the polygenic score. For example, in the most recently published GWAS the variance explained for educational polygenic scores on phenotypic educational attainment was 12% (6). In the same study, polygenic scores for cognitive performance explained 7-10% of the variance in cognitive performance. Further, based on case-control differences, the polygenic score for schizophrenia (8), MDD (9), and anxiety (55) explained at most 7%, 1.9%, 2.3% and 2.1% of variance in liability, respectively. The low predictive accuracy of the polygenic risk scores may partly be due to the heterogeneity of complex human traits and disorders, which may disguise true genetic pleiotropy between brain features and the polygenic architecture of complex traits. Dimensional approaches such as normative modeling (56, 57) or brain age prediction (58) may reveal stronger shared signal. Many studies have shown that complex traits and disorders have substantial genetic overlap (59, 60) which may dilute the signal-to-noise ratio and specificity of polygenic scores. Future studies applying GWAS results with more power and novel techniques for boosting the shared genetic signal between traits (61, 62) may reveal stronger evidence of pleiotropy between brain FC and the polygenic architecture of complex traits. Taken together with other inherent limitations (see Supplements), the converging pattern across metrics of model performance indicate chance-level performance of brain FC for polygenic scores.

In conclusion, based on fMRI data from 10,343 individuals we have demonstrated high prediction accuracy of fMRI-based brain connectivity for age and sex, as well as above chance accuracy for educational attainment and intelligence (cognitive). In contrast, we obtained chance-level prediction accuracy for dimensional measures of depression, anxiety and neuroticism (mental health), in addition to their genetic underpinnings based on the respective polygenic scores and polygenic risk for schizophrenia. These novel findings support the link between cognitive abilities and the brain organization of the brain functional connectome and provide a reference for imaging studies mapping the brain mechanisms of clinical and genetic risk for mental disorders.

## Materials and Methods

### Sample

We used the October 2018 release of UKB imaging data (63), comprising 12,213 individuals with resting-state fMRI scans that passed the quality control performed by the UKB (64). The study was funded by the Research Council of Norway, the South-Eastern Norway Regional Health Authority, and approved by the UKB Access Committee (Project No. 27412) and the South-Eastern Norway Regional Committees for Medical and Health Research Ethics (REC). All participants provided informed consent prior to enrolment. The exclusion criteria across all phenotypes and polygenic scores were individuals with an ICD-10 based mental or a neurological disorder (N = 210) and non-Caucasians (N = 1659) yielding a total of 10,343 individuals. Most of these participants were MRI-scanned in Manchester, but some were MRI-scanned in Newcastle (N = 354).

### Phenotype data

Neuroticism was calculated based on a previous implementation (65), while educational attainment was based on the “qualifications” variable (UKB field: 6138) following a recent imputation procedure (66). Fluid intelligence (UKB field: 20016) was defined as the sum of the number of correct answers on the verbal-numerical reasoning test (13 items). Data for all the aforementioned phenotypes were taken from instance 2 (imaging visit). Symptom load for depression and anxiety were based on the sum score from the 9-item Patient Health Questionnaire (PHQ-9) and the 7-item Generalized Anxiety Disorder (GAD-7) questionnaire, respectively. Both PHQ-9 and GAD-7 were taken from the online follow-up. Table S2 shows the specific number of individuals for each phenotype and all polygenic scores.

### Polygenic risk scores

Procedures for DNA collection and genotyping in UKB have been described previously (67). Polygenic scores were calculated using PRSice v. 1.25 (68) based on GWAS results for broad depression, probable MDD (69), diagnostic MDD (9), item-level and sum neuroticism (65), anxiety (70), and schizophrenia (8). In the case of polygenic scores based on item-level and sum neuroticism, we performed our own GWAS (see Supplemental Material) to avoid overlap in participants in the discovery and target dataset (i.e. fMRI sample). We used the same reasoning to run our own GWAS (see Supplemental) to derive polygenic scores for educational attainment (based on years of schooling) and fluid intelligence, following similar procedures to previous studies (6, 7). For the main polygenic score analyses, a liberal SNP inclusion threshold was set at p ≤ 0.5, as this typically explains more variance in the clinical phenotype (71). For sensitivity analyses, we repeated the same analyses using a more conservative threshold of p ≤ 0.05. As an additional sensitivity analysis, we ran a PCA on the computed polygenic scores from thresholds p ≤ 0.001 to p ≤ 0.5 (at 0.001 intervals) based on a recent implementation (36).

### Image Acquisition and pre-processing

Detailed description of the image acquisition, pre-processing, group-level ICA and dual regression can be found in a previous study (64). The majority of the MRI data used were obtained in Cheadle Manchester on a Siemens Skyra 3.0 T scanner (Siemens Medical Solutions, Germany) with a 32-channel head coil. A small number of scans (N = 354) were obtained at an identical scanner in Newcastle. FSL (http://fsl.fmrib.ox.ac.uk/fsl) was used for fMRI data preprocessing. Briefly this involved motion correction, high-pass temporal filtering, echo-planar image unwarping, gradient distortion correction unwarping, and removal of structured artifacts. Estimated mean relative in-scanner head motion (volume-to-volume displacement) was computed with MCFLIRT. Group-level ICA was carried using MELODIC based on 4,162 datasets with a dimensionality of 25. Four of these ICs were identified as noise and discarded, leaving a total of 21 ICs for analyses. Dual regression was performed to generate subject specific spatial maps and corresponding time series.

### Functional connectivity measures

All FC measures were computed in MATLAB. SFC was computed both with the unfiltered and bandpass filtered time-series within 0.04-0.07 Hz. For both, a node-by-node connectivity matrix was created using partial correlations between the subsequent time-series (72), resulting in 210 unique edges. These partial correlations were L1-regularized, with estimated regularization strength (lambda) at the subject level (73–75).

For dFC we used a phase-based method (76) within the 0.04-0.07 Hz frequency band (77). Briefly, this method is sensitive to the degree of coupling and de-coupling between pairs of brain networks across the scanning session, based on applying the Hilbert transform and subsequently the Kuramoto order on the node time-series.

### Statistical analysis

All statistical analyses were performed in R version 3.4.2 (78). We z-normalized all polygenic scores prior to the multivariate analyses.

### Multivariate analysis of brain FC

In our primary analyses, we tested to what extent various combinations of static and dynamic FC between all nodes in the an extended brain network could predict the phenotypes, by using shrinkage linear regression (79) implemented in the R-package ‘care’ (http://strimmerlab.org/software/care). See the original paper for details on the optimization of shrinkage parameters. Firstly, 80% of the data was used as the training set while the remaining 20% of the data was used as the left-out test set. We ran 10-fold internal cross-validation on the training set (i.e. based on iteratively using 90% of the sample to predict the remaining 10%), repeated 100 times on randomly partitioned data. We computed RMSE as our main measure of model performance for all phenotypes, but also MAE and R^2^ (explained variance or model fit). R^2^ was mainly used to visualize and compare model performance across models. We also used spearman’s *rho* as a measure for model performance for predicting these phenotypes. For all phenotypes, we computed Spearman’s rho, but Pearson correlation coefficient (*r*) for age. For sex, we used logistic regression for classification and accuracy, sensitivity and specificity as markers of model performance. The same model parameters were used in the model validation, by using the whole training set to predict the phenotypes in the left-out test set. Statistical significance for phenotypes were assessed for the best feature set with a positive R^2^ using permutation-based testing (10,000 permutations) based on RMSE, with an additional Bonferroni correction for 7 phenotypes and 20 polygenic score models. The mean model performance from cross-validation of the empirical data was used as our point estimate. For the phenotype models that were statistically significant, we determined the relative importance of each edge by computing the mean CAR scores (80) in cross-validation. We also assessed the correlation amongst these models based on the CAR-scores. To assess the confounding effects of age, sex and head motion, we repeated the analyses for fluid intelligence by regressing these confounders from the edges in the sFC feature set. We used the same framework to predict polygenic scores, computing Pearson’s r in addition to the other measures of model performance, and model validation for the main PGRS analyses (p ≤ 0.5). To assess the effect of population stratification, we regressed out the first 10 genetic principal components genetic principal components from the edges of the best feature set for one of the polygenic score models. We also assessed the correlation amongst phenotypes and corresponding polygenic scores based on the CAR-scores in the cases where the phenotype model was statistically significant.

## Acknowledgments

The authors were funded by the Research Council of Norway (213837, 229129, 204966, 249795, 251134), the South-Eastern Norway Regional Health Authority (2014097, 2015073, 2016083, 2017112), the Department of Psychology, University of Oslo and the KG Jebsen Stiftelsen (223723). This research has been conducted using the UK Biobank Resource (access code 27412). The permutation testing was performed using resources provided by UNINETT Sigma2 - the National Infrastructure for High Performance Computing and Data Storage in Norway.

