## Supplementary material for "Predicting cognitive and mental health traits and their polygenic architecture using large-scale brain connectomics": Supplmental Information

**Supporting Information**

**Methods**

*GWAS without imaging subset*

To prevent overlap between the base and target samples of the polygenic scoring, we ran a GWAS on the UK Biobank (UKB) cohort excluding the individuals with neuroimaging data available. For this, we used PLINK 2.0 and the UKB October 2018 release imputed genetic data, filtering out SNPs with a minor allele frequency below 0.001 or failing the Hardy-Weinberg equilibrium test at p < 1x10-9. We further removed all non-Caucasians and individuals with a brain disorder as indicated by ICD10. As such, the final sample size for these GWAS was n = 402,515. We ran a logistic regression GWAS for the individual neuroticism items (absent/present), and a linear regression for the neuroticism sum score, educational attainment and fluid intelligence, covarying for age, sex and ten genetic principal components. The neuroticism items are from the Eysenck Personality Questionnaire-Revised Short Form, and covered the following behaviour domains: mood (UKB field: 1920), miserable (UKB field: 1930), irritable (UKB field: 1940), feelings easily hurt (UKB field: 1950), fed up (UKB field: 1960), nervous feeling (UKB field: 1970), worry (UKB field: 1980), tense (UKB field: 1990), worry of embarrassment (UKB field: 2000), suffering from nerves (UKB field: 2010), loneliness (UKB field: 2020) and guilt (UKB field: 2030)

*Hierarchical clustering of resting state networks*

The brain nodes from ICA were grouped together using hierarchical clustering (see Supplemental Figure 12). Briefly, the node-by-node connectivity matrix based on average full correlation (fisher-z transformed) were submitted to hierarchical clustering as implemented in FSLNets (1), based on the Euclidean distance and ward linkage function in MATLAB 2014B (The Mathworks Inc.).

### Results

*Connectome overlap of traits and polygenic scores*

The correlation between phenotypes for specific feature sets based on CAR-scores that had a mean model performance with a postive $R^2$ value were as follows: fluid intelligence and educational attainment (sFC = 0.612, bandpass filtered sFC = 0.520, dFC = 0.457), age and intelligence (sFC = -0.240, bandpass filtered sFC = -0.349, dFC = -0.178), age and education (sFC = -0.217, bandpass filtered sFC = -0.201, dFC = -0.234). The correlation between the CAR scores obtained from the models predicting a trait and its corresponding polygenic scores were overall positive: educational attainment (sFC = 0.618, bandpass filtered sFC = 0.549, dFC = 0.449), fluid intelligence (sFC = 0.504, bandpass filtered sFC = 0.381, dFC = 0.403).

### Limitations

Years of educational attainment were converted from educational qualification, meaning that we are not sampling the true years of educational attainment. Furthermore, this variable is skewed, with an over-representation of highly educated individuals. The verbal-numerical test is purportedly a poor measure of fluid intelligence, partly considering that it  only consists of a limited number of items based on a correct or incorrect response. The dimensional measures of depression anxiety and neuroticism were significantly positively skewed, given that all the individuals are generally healthy, which likely affects prediction accuracy. Both the measures of depression and anxiety were only available from the online-follow up and not the imaging visit, meaning that there is substantial temporal lag.

A few studies have indicated that the number of nodes (e.g. dimensionality of the ICA decomposition) influences prediction accuracy, with an estimated optimal number ranging from 50 (2) to around 150 (3). Caution is warranted with regards to over-interpretation of the edge-wise feature importance, as these are interpretable only in the context of the specific multivariate model. Despite this, we see a similar pattern of feature importance of sFC-edges in the feature set with only sFC edges and the feature set with all three FC types. Relatedly, it is possible that different definitions of brain regions (e.g. seed-based) would yield different results. Despite this, in terms of prediction accuracy, a study found evidence that nodes defined using information in the data (e.g. ICA) gave the best model performance (3).

Another limitation in the current study is that the polygenic scores for broad and probable depression were derived from the same dataset that the GWAS was based on, which likely leads to overfitting. However, model performance for these polygenic scores were low, and thus do not affect interpretation of the results. Unfortunately, we did not have a (dimensional) phenotypic counterpart for the schizophrenia polygenic score. Caution is warranted when interpreting polygenic scores of educational attainment and fluid intelligence from current GWAS, as a preliminary study found that they are confounded by socioeconomic effects (4).

### Tables

**Table S1**: The correlation (Pearson's r) between the feature importance of the different FC-types (sFC, bandpass filtered sFC and dFC) within phenotypes

| | Age | Fluid intelligence | Educational attainment |
|---|---|---|---|
| sFC – sFC band | -0.198 | -0.181 | -0.104 |
| sFC – dFC | -0.741 | -0.647 | -0.677 |
| sFC_band - dFC | 0.157 | 0.042 | 0.042 |

**Table S1**: The number of individuals with data for each variable used in the multivariate machine learning analyses.

| **Variable** | **Number of individuals** |
|---|---|
| Age | 10,343 |
| Sex | 10,343 |
| Educational attainment | 10,251 |
| Fluid Intelligence | 9,607 |
| Sum neuroticism | 10,115 |
| Sum depression symptoms | 7,777 |
| Sum anxiety symptoms | 7,815 |
| All polygenic scores | 10,320 |

### Figures

### Multivariate brain associations

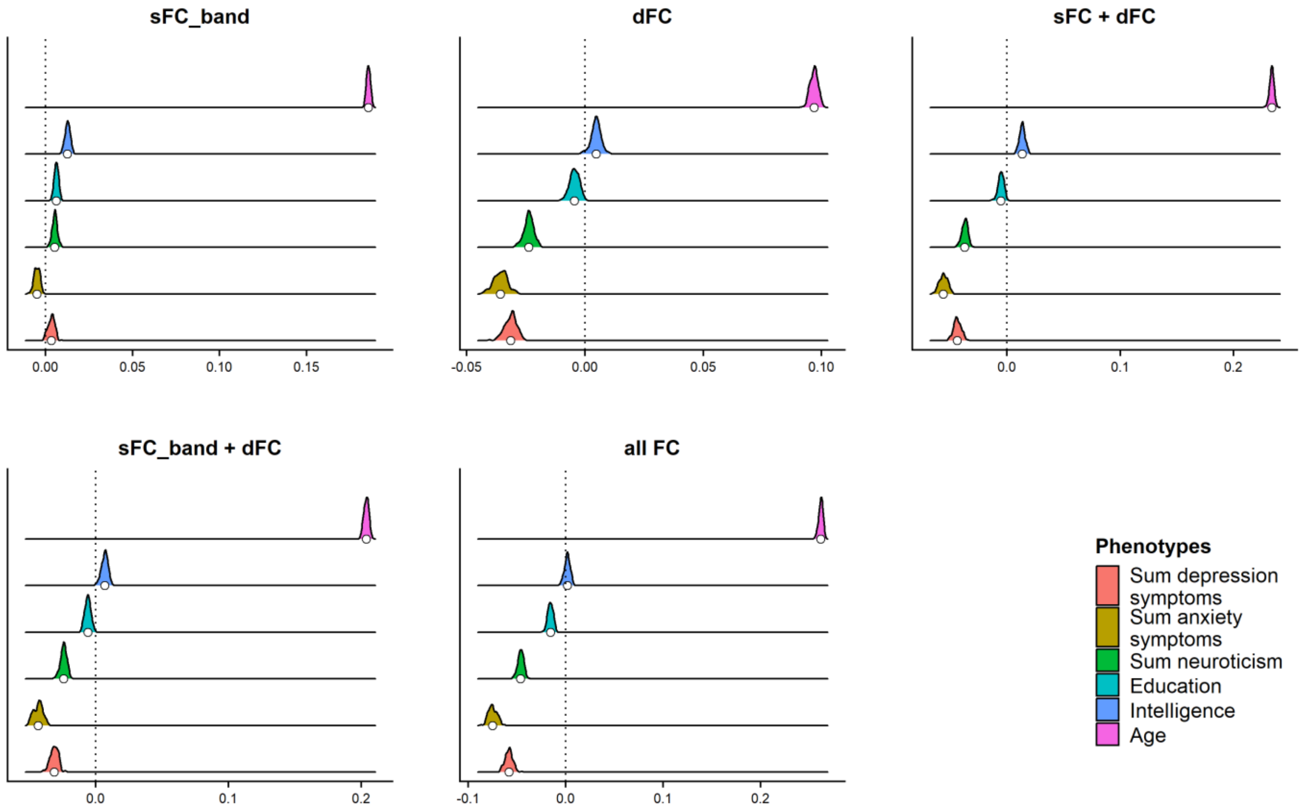


**Fig. S1.** A: The bars represent the cross-validated results of each FC feature set for age, educational attainment, and intelligence. The y-axis represents the Pearson's correlation between raw and predicted age as a marker of model performance. For sum neuroticism, educational attainment, and intelligence, spearman's rho is the model performance. Model performance based on $R^2$ for sum neuroticism was slightly above zero using the filtered sFC feature set, but not statistically significant based on RMSE (permutations testing with additional Bonferroni correction).

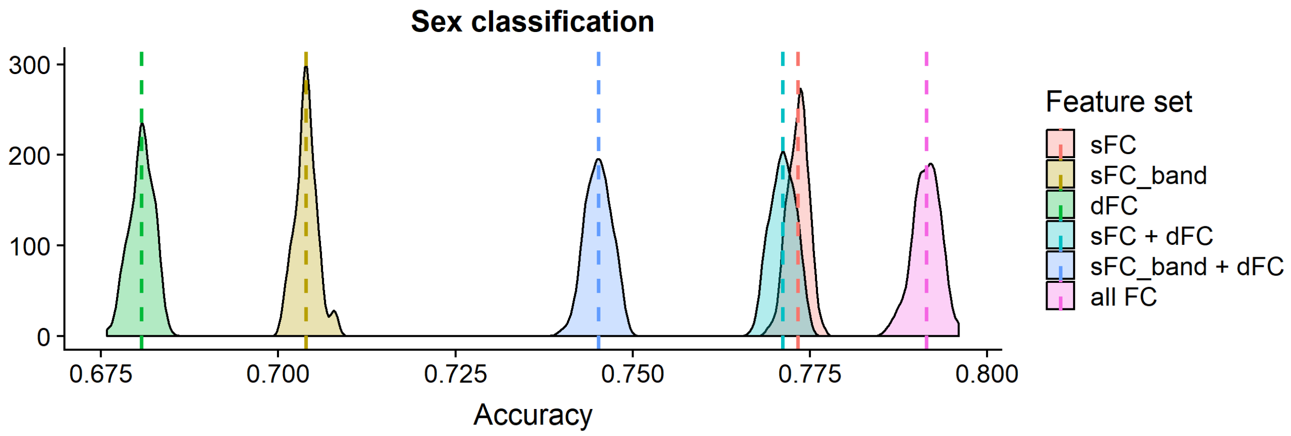


**Fig. S2.** Cross-validated results of all the feature sets for sex classification. The dashed vertical lines represent the mean accuracy for a given feature set.

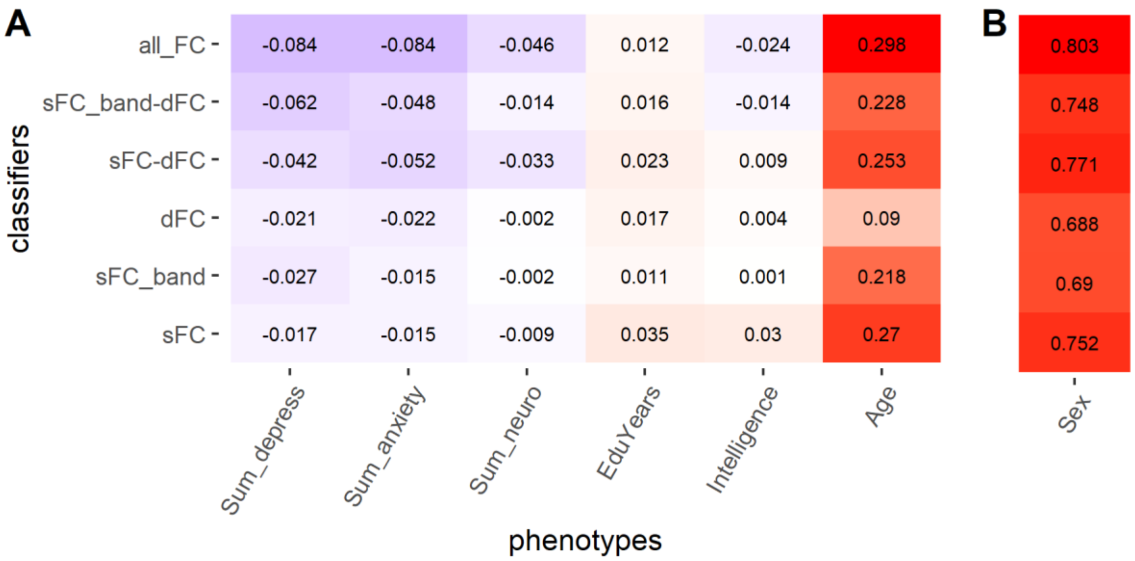


**Fig. S3.** Final model validation results predicting (A) age, fluid intelligence, educational attainment, and dimensional measures of depression, anxiety and neuroticism ($R^2$), and (B) sex (accuracy) for different combinations of the FC feature sets.

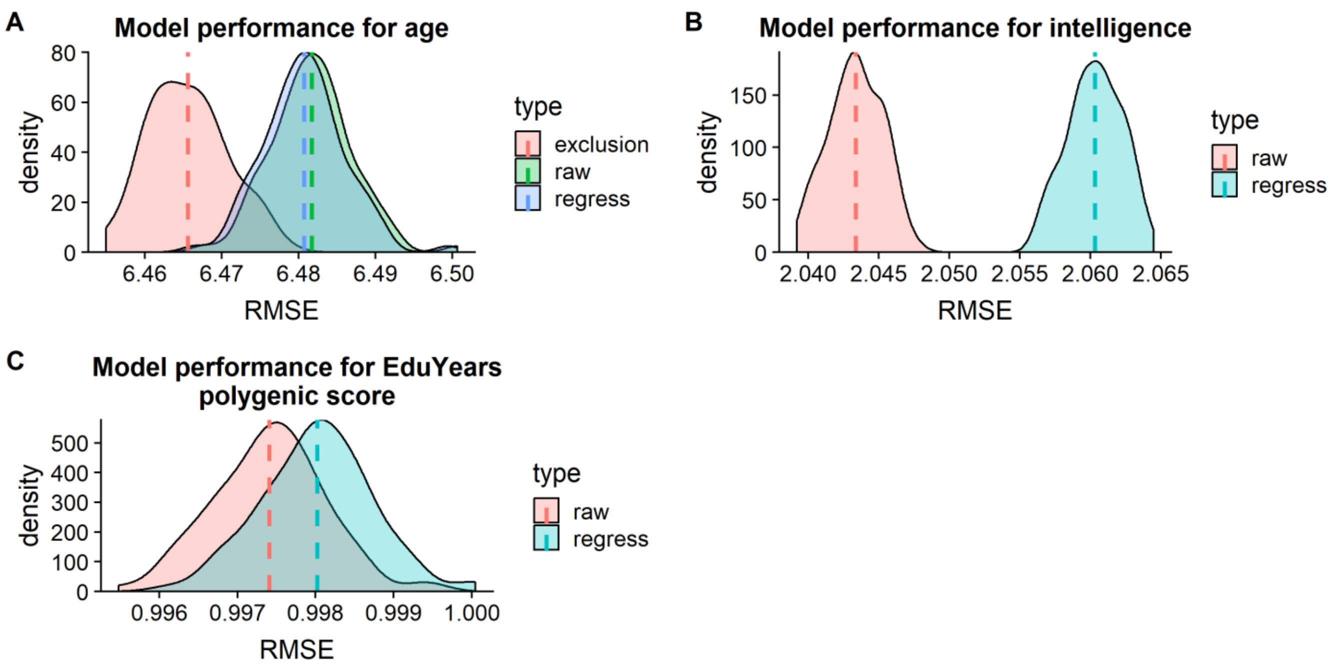


**Fig. S4** Comparing results that have and have not been accounted for various confounders in the sFC feature set. (A) Predicting age without addressing scanner site (raw), regressing out scanner site from the FC edges (regress) and by excluding participants from Newcastle-Upon-Tyne (exclusion). (B) Predicting intelligence by not addressing confounds (raw) and regressing out sex, head motion and linear and quadratic effects of age from the edges. (C) Predicting educational attainment polygenic score without addressing confounds (raw) and addressing population stratification by regressing out the first 10 genetic principal components from the edges in the sFC feature set (regress).

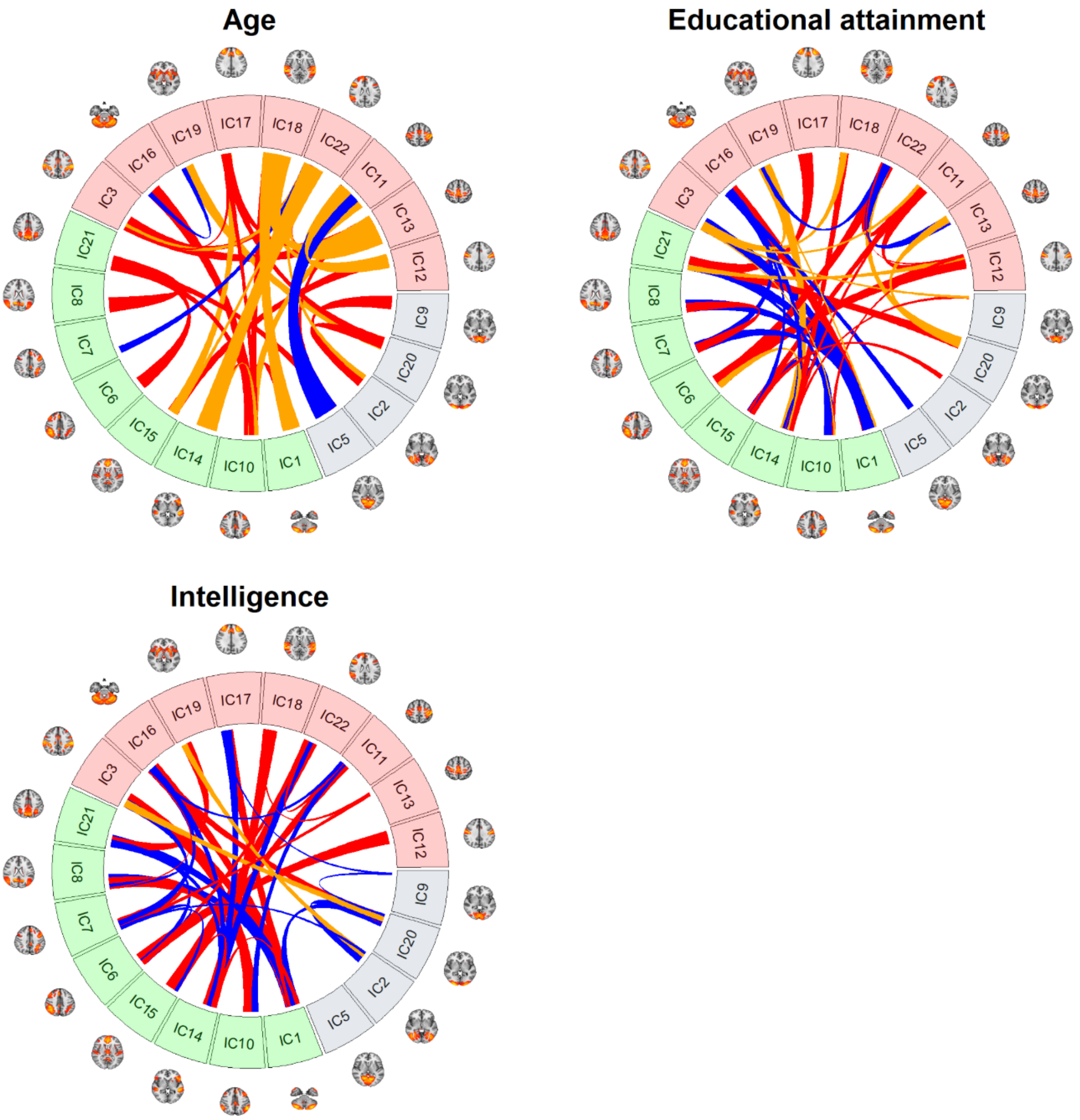


**Fig. S5.** The top 40 edges based on CAR-scores for the feature set that includes sFC (red), bandpass filtered sFC (orange) and dFC (blue). Cross-fold Validation for the age, phenotypic years of educational attainment and fluid intelligence. These represent the relative importance of each edge positively associated with the specific trait, with the thickness of the line signifying the relative order.

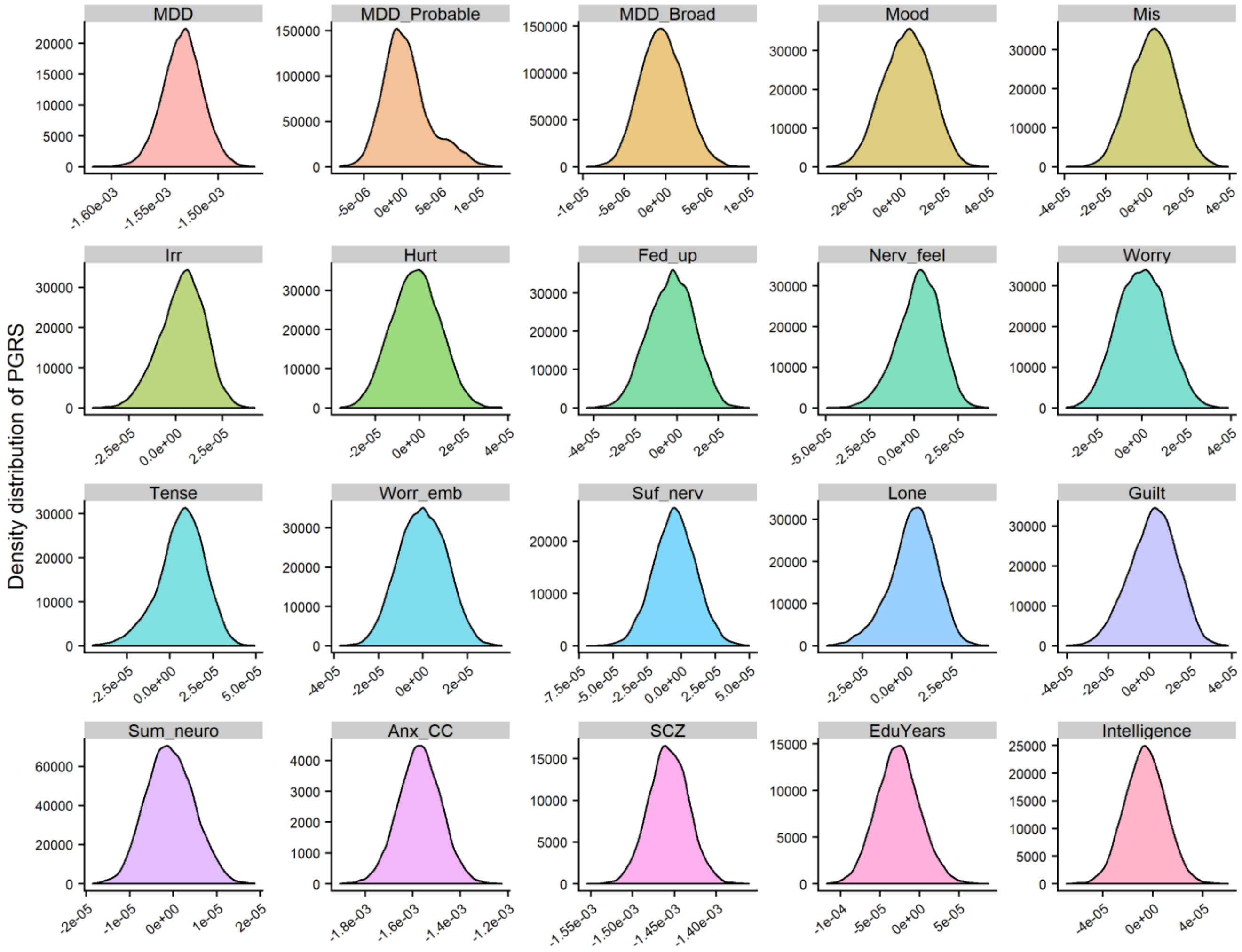


**Fig. S6.** Distribution of each polygenic scores (raw values)

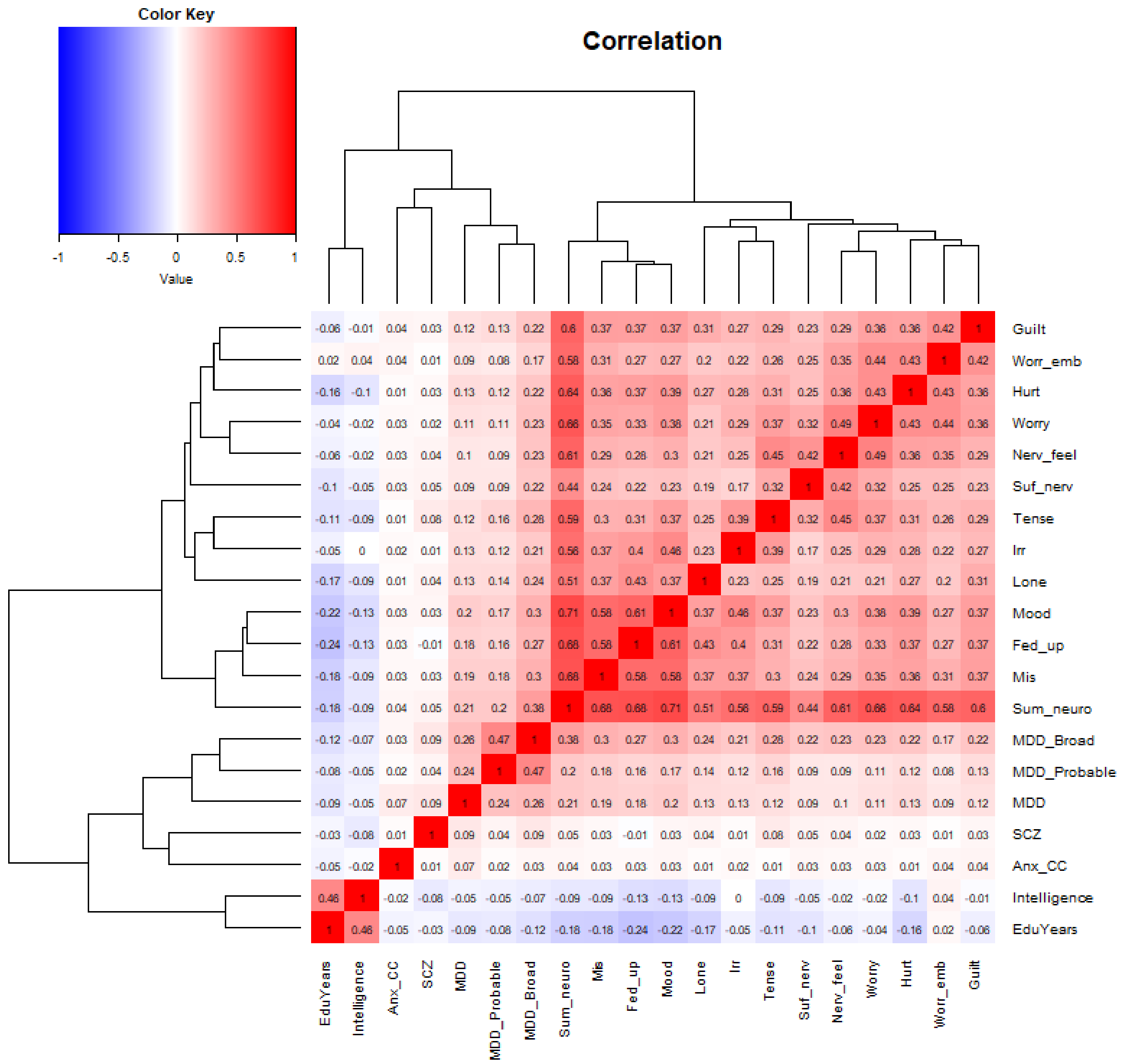


**Fig. S7.** Heatmap of the correlation between all the (raw) polygenic scores (with threshold p ≤ 0.5). The polygenic scores are ordered in terms of hierarchical clustering as indicated by the dendrograms

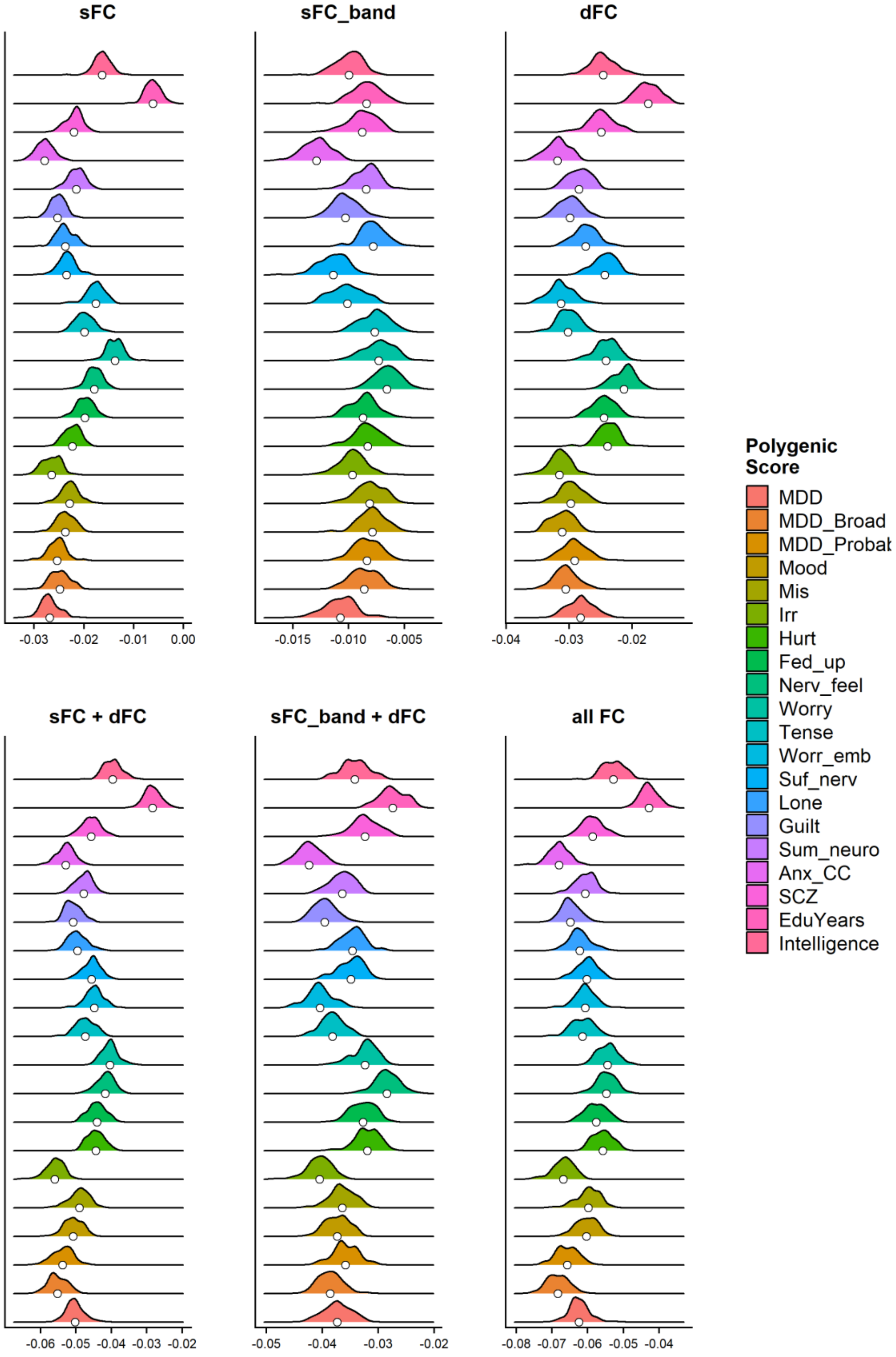


**Fig. S8.** Cross-validated results based on $R^2$ for the various FC feature sets for the various polygenic scores

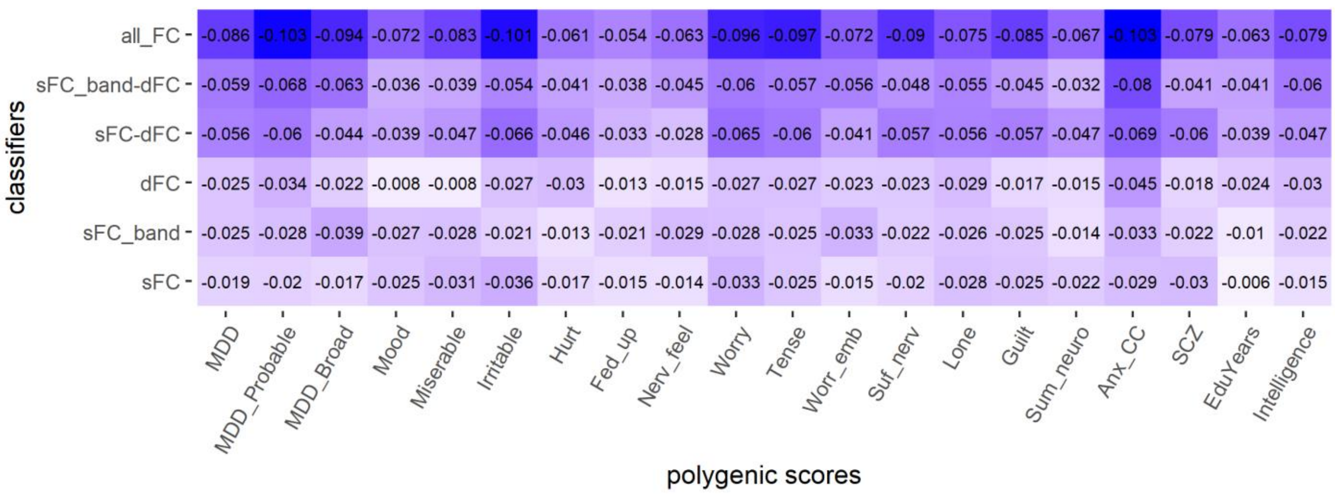


**Fig. S9.** Final model validation results ($R^2$), predicting polygenic scores for different combinations of FC feature sets.

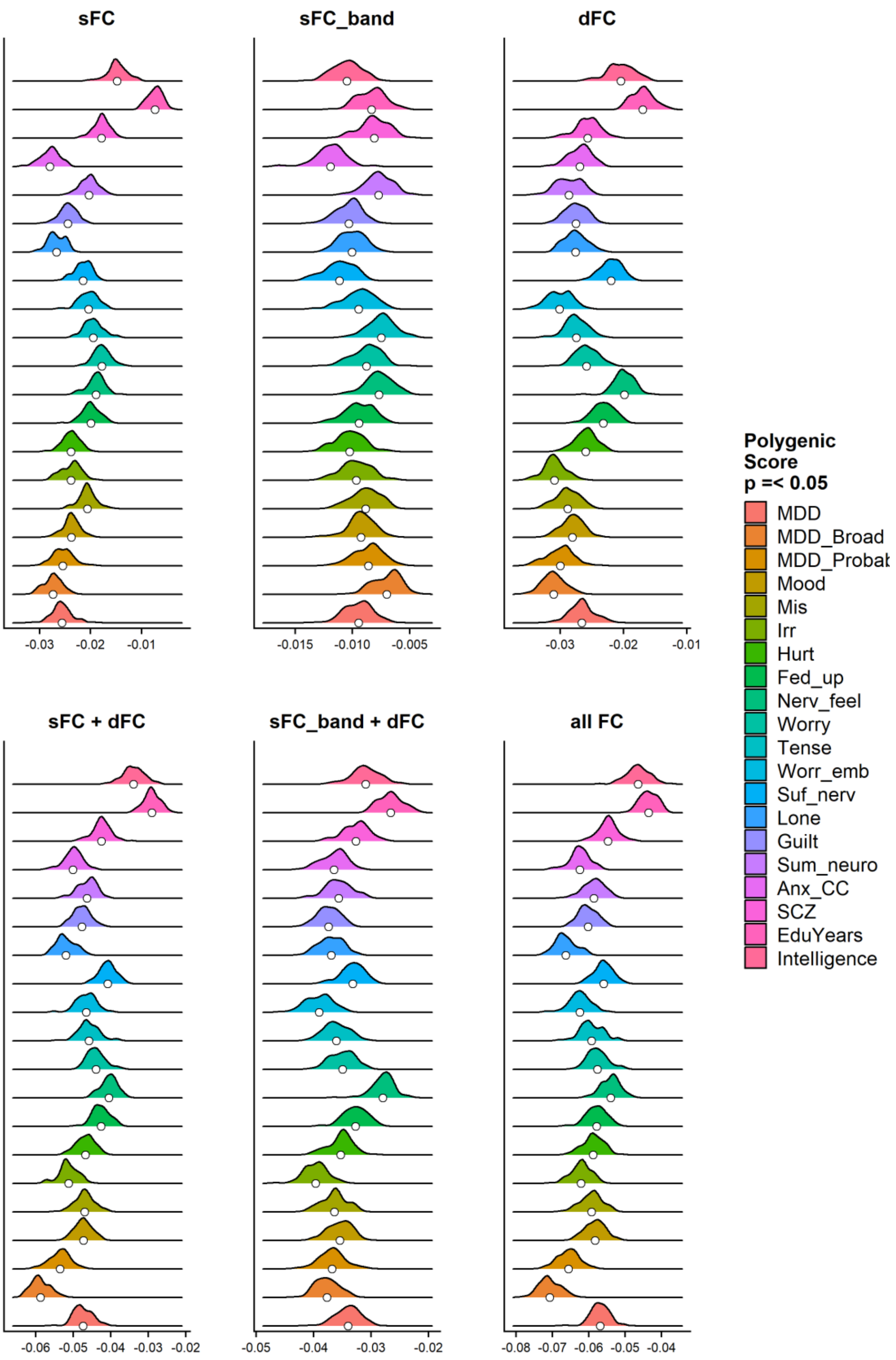


**Fig. S10.** Cross-validated results, predicting polygenic scores thresholded at $p \leq 0.05$ for different combinations of FC feature sets.

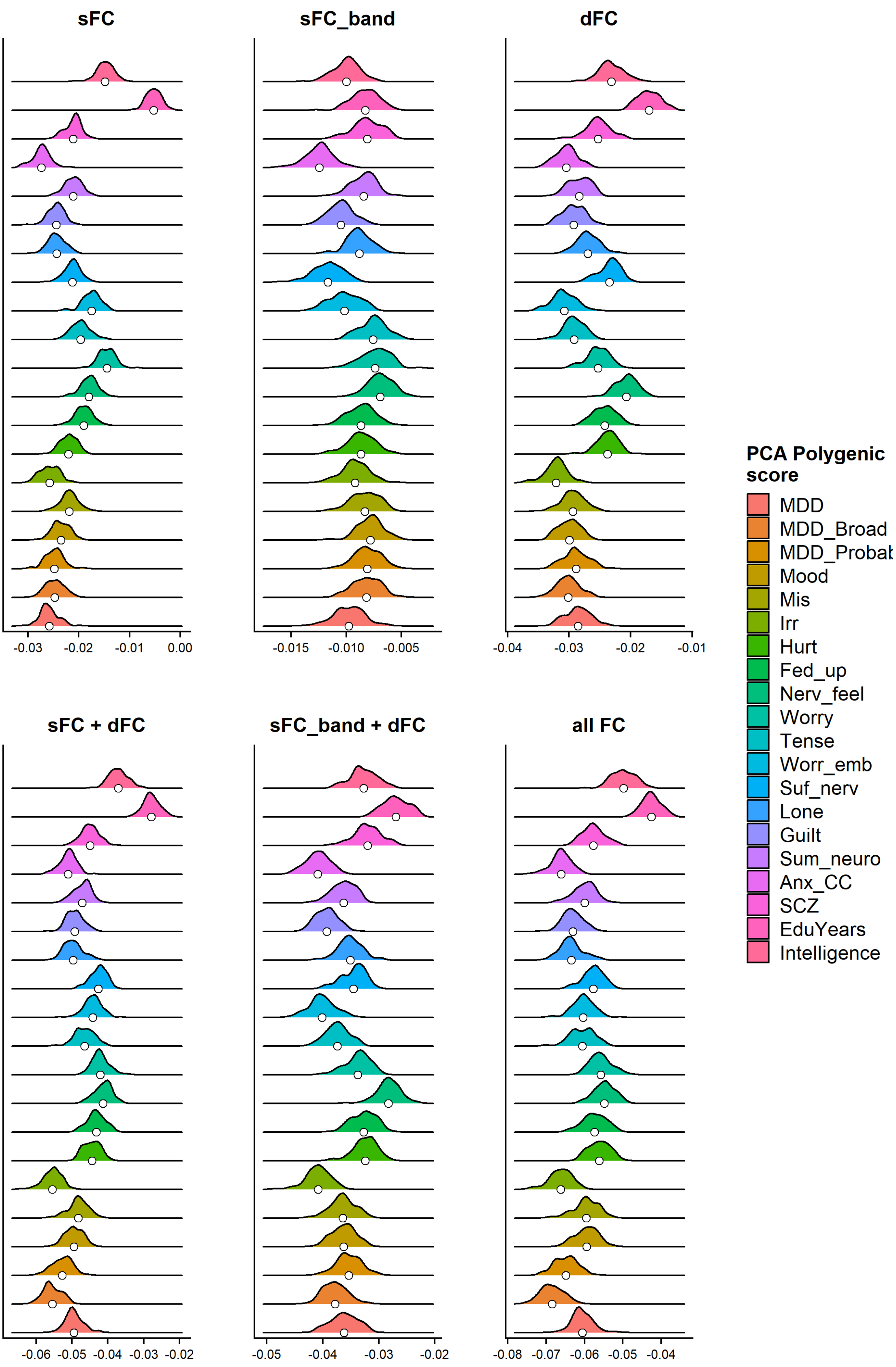


**Fig. S11.** Cross-validated results, predicting the first component of PCA-polygenic scores for different combinations of FC feature sets.

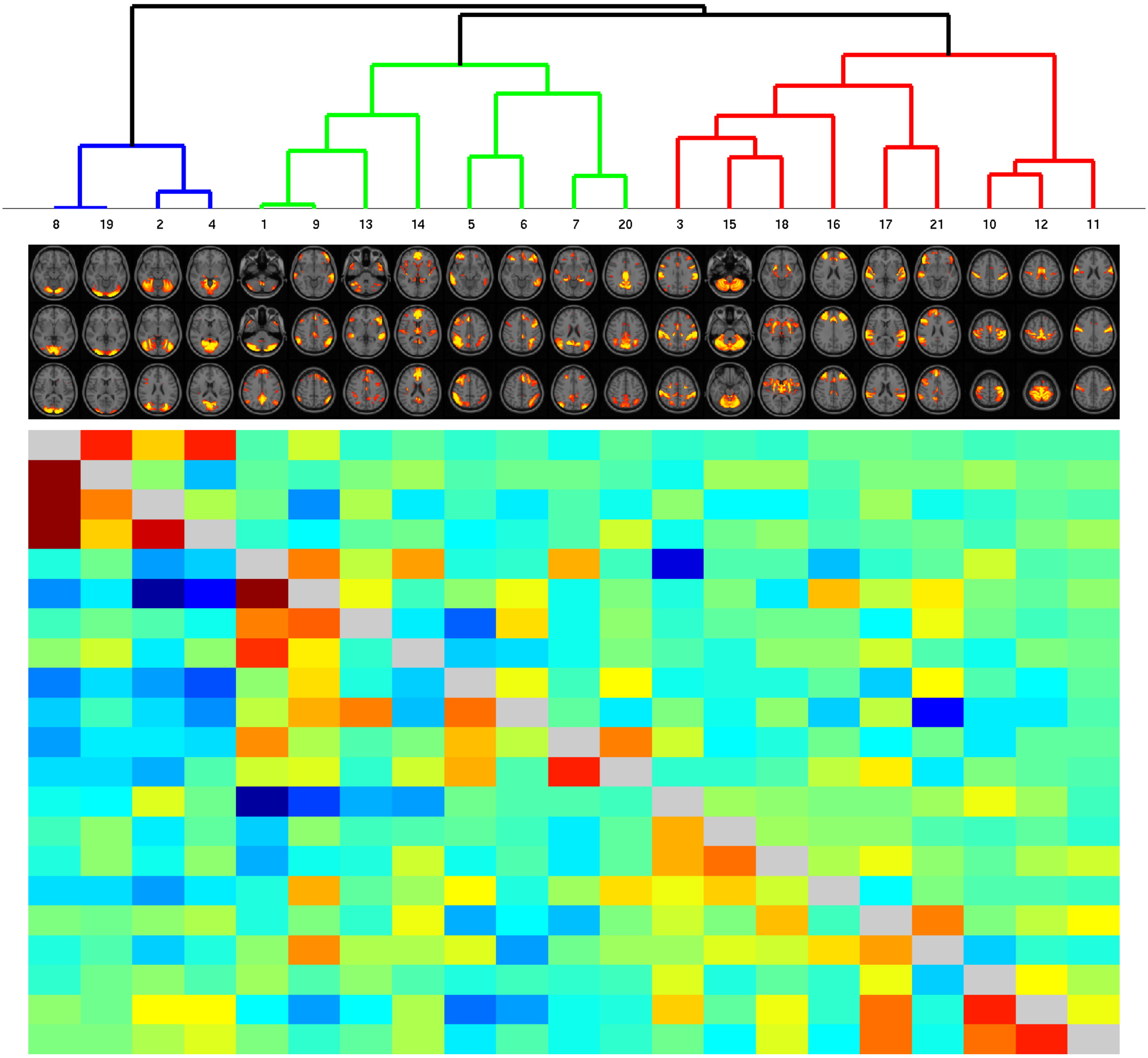


**Fig. S12.** Hierarchical clustering of all the included brain components from ICA based on the full correlation between IC nodes of all subjects. Each IC node is denoted by one column. The lower diagonal matrix displays the full correlation of the time-series between components. Dark red indicates a high positive correlation, green indicates 0 correlation, and dark blue represents high negative correlation. The upper diagonal matrix displays the regularized partial correlation of the time-series between components.
